# Agent-based simulation for reconstructing social structure by observing collective movements with special reference to single-file movement

**DOI:** 10.1101/2020.03.25.007500

**Authors:** Hiroki Koda, Zin Arai, Ikki Matsuda

## Abstract

Understanding social organization is fundamental for the analysis of animal societies. In this study, animal single-file movement data ‒serialized order movements generated by simple bottom-up rules of collective movements— are informative and effective observations for the reconstruction of animal social structures using agent-based models. For simulation, artificial 2-dimensional spatial distributions were prepared with the simple assumption of clustered structures of a group. Animals in the group are either independent or dependent agents. Independent agents distribute spatially independently each one another, while dependent agents distribute depending on the distribution of independent agents. Artificial agent spatial distributions aim to represent clustered structures of agent locations ‒a coupling of “core” or “keystone” subjects and “subordinate” or “follower” subjects. Collective movements were simulated following two simple rules, 1) initiators of the movement are randomly chosen, and 2) the next moving agent is always the nearest neighbor of the last moving agents, generating “single-file movement” data. Finally, social networks were visualized, and clustered structures reconstructed using a recent major social network analysis (SNA) algorithm, the Louvain algorithm, for rapid unfolding of communities in large networks. Simulations revealed possible reconstruction of clustered social structures using relatively minor observations of single-file movement, suggesting possible application of single-file movement observations for SNA use in field investigations of wild animals.

## Introduction

Identifying latent structures underlying animal social organization is fundamental in animal ecology [1], as well as in the social sciences [2]. In order to infer the biological mechanisms underlying the emergence of complex animal societies, researchers have traditionally recorded animal social data in the multiple dimensions of animal societies; for example, spatial distances among group members, social relationships (male–female or mother–offspring relationships), and movements based on group decision making.

The recent update of computational models for social network analysis (SNA) enables quantitative evaluation of complex processes of social networks based using big data. The basic framework of SNA envisions multiple nodes as animals, viruses, people, or any other agents within the network and at its edges as a dyadic relation mathematically defined between the two nodes (e.g., [3–5]). The SNA aims to statistically link network structures with bottom-up rules defining dyadic relations, e.g., affiliations, friendships, proximities, co-occurrences, follower/followee, signal sender/receiver, or communications [6]. SNA provides a modern powerful interface for visualizing complex biological networks.

One major biological interest for SNA is statistical approximations of modular, clustered or hierarchical structures embedded in animal social groups [1, 7]. Doubtless, kinship is a basic concept in the theory of the socioecology (for a classical kickoff, e.g., [8]). A mother invests effort in her offspring, mother and offspring generally maintain proximity, and degrees of the social bonding are reflected by relatedness [9, 10]. Based on the cumulative effects of such relatedness, social structures are determined by additional social relations, such as male-female sexual interactions and male-male competition [11, 12]. Consequently, a wide variety of social roles in the animal group are recognized, and multi-layered clustered structures of the group emerge [1, 13–15]. Some metrics commonly implemented in the modern SNAs provide statistical indices for modularity, centrality or cohesiveness (e.g., [16]), and have been applied to actual observational data of animals.

For the practical applications of SNA to observational data, dyadic social interactive behaviors or distance-based proximities have been commonly used in ecological analyses of animal societies [3, 17]. Allogrooming is a typical example of dyadic social interaction that is widely observed in social mammals. Allogrooming is an altruistic behavior and an appropriate behavioral metric for evaluating affiliative interactions (e.g., [18–20]). Likewise, agonistic interactions, where dominant animals show aggression toward subordinate animals, are also interpretable and countable behaviors that can be used to characterize dominance hierarchies (e.g., [21–23]). Generally, in the case of dyadic social interactions, researchers calculate the occurrence frequencies of dyadic events using an observational rule; for example, counting the grooming occurrences and recording the identity of the groomer and groomee in a specific observational session. These data obtained for social interactions can be used to construct matrices of social relationship strength by scoring or weighting the obtained data of dyadic relationships [17]. However, despite the interpretable advantages of social relationships, there are certain disadvantages associated with such data. Behavioral-based metrics such as grooming or fighting are generally unsuitable for the automation of data acquisition or evaluation. Currently, at least, researchers must record animal behaviors via direct observation or video recording for data analysis. Additional disadvantages are the “specializations” of social behaviors; for example, grooming might not always be observed in insects, or we may have little knowledge regarding the agonistic interactions among some animals, thereby limiting the analysis. In contrast, distance-based data lack contextual information and are not perfectly linked to social relationships [24]. In the case of distance-based proximities, inter-individual distances are recorded using a regular cycle of sampling rates (e.g., recording the distance at 5-min intervals), and proximities are evaluated. Subsequently, a matrix of social relationships is generated in a similar manner, scoring or weighting the proximity. Given that close proximity is assumed to indicate an affiliative relationship, the thresholds of the specific distance defining the social “association” can be used to generate binary data (i.e., association or isolation), and an adjacency matrix is generated from a summation of the binary data [17]. These data are generally accepted to indirectly reflect social affiliations [25]. Affiliated individuals may maintain proximity as typically seen in mother-offspring pairs. The recent advanced tracking methods using high-precision GPS (e.g., [26, 27]) or Bluetooth beacons (e.g., [28]) have strongly influenced location-based proximity metrics due to dramatic expansion of data for interactions among “trackable” animals. Real-time distance-based proximities among all animal agents empirically reveal core mechanisms underlying collective movement emergencies. Simultaneous tracking of multiple animals using wearable tags is technologically possible. However, perfect tracking of suitable numbers of animals in groups/units for analyses of collective movement are still limited to laboratory-based animals (e.g., [28, 29]) or capturable wild animals [30–32]. Thus, a key analytical step for SNA is selection of appropriate data, which researchers record massively, easily, reliably and precisely.

The main objectives of this study were (1) to propose a novel approach for obtaining observable data that has not been systematically applied in the estimation of animal social organization, and (2) to provide methodological validation for the proposed approach. The first objective is based on practicality, particularly with respect to those studying wild animals. Dyadic social interactive behaviors or distance-based proximities have been a golden standard used to infer animal social organizations, as mentioned above; however, those studying animal behavior often experience difficulty in obtaining large amounts of high-quality data for such dyadic social interactions or spatial proximity in wild animals. For example, it is extremely difficult to monitor complete behavioral activities, even if wearable sensors are available. Similarly, in most cases, experimental manipulations, such as artificial control of specific individual movements, are not possible for wild animals (for a discussion of the methodological issues, see [17]). Accordingly, researchers invariably search for alternative promising types of data, together with technological updates. For the generalized application of novel types of data, empirical evidence that can be used to validate the approaches is necessary, which could partly be achieved using numerical simulations.

In this study, another kind of testable data for the SNA is proposed, that is, “animal single-file movement” generated by bottom-up rules in the theory of collective movements. This topic is currently of great interest in animal ecology (for a recent review, e.g., [33]). Typically in fish schools, bird flocks, insect swarms, or ungulate herds, large numbers of individual agents show highly-coordinated collective movements [34, 35]. Theoretically, collective movements do not rely on decision-making of specific individuals, e.g., a “leader”, but emerge in a bottom-up manner such as iterated interactions between animals, either directly (e.g. via visual cues) or indirectly (e.g. via trail formation).

Animal single-file movement, defined as serially ordered patterns of animals that are indirectly determined by degrees of inter-agent relations, is a promising target for data collection for linking SNA linking with recent discussions of collective movement by social animals. First, single-file movements are observable in many free-ranging animal species. Classically, social animals moving in single file provided an excellent opportunity to count group members inhabiting dense forests with poor visibility, and further to infer social structures. Extensive historical fieldwork reveals that such events provide information on latent key factors underlying animal social organizations, e.g., dominance hierarchy [36, 37], kin relationships [38], and physiological status, such as breeding cycle [39, 40]. Additionally, due to the recent human imposed threats such as roads, animals are often forced into single-file formation to cross risky or dangerous places. Such an aspect of socio-spatial organization under highly risky situations has also been analyzed in meerkat [41] and elephant [42]. In fact, recent studies focusing on single-file movement and social system structure have examined several animal taxa, e.g., wolves, *Canis lupus*: [36]; zebra, *Equus burchellii*: [39], buffalo, *Syncems cuffer*: [40], chimpanzee, *Pan troglodytes*: [43], mandrill, *Mandrillus sphinx*: [44] and spider monkey, *Ateles geoffroyi*: [45]. Second, animal single-file movement can be recorded without capture and release for attachment of location tags, simply by video-recording (e.g., camera-trappings). This advantage for studying collective movement avoids use of GPS collars that are a burden for free-ranging animals, and most cannot be used for threatened and endangered species [46]. Also, single-file movement is naturally observed even in zebra heading to water holes [39] or in wolves through snow [47]. Anywhere such single-file movements are often observable recently developed camera-trapping techniques [48] would enable collection of a valuable data set. Further, where animal movement is controllable in semi-free range conditions such as narrow gateways to feeders and water, another opportunity exists. Such single-file movement has been recorded for sheep, *Ovis aries*: [37]. Third, serial order patterns governed by collective movements allow behavioral ecologists to examine individual-based observation essential for documentation and measurement of animal behavior. Fourth, the methodological advantages listed above compensate for the disadvantages of other metrics commonly used in the previous studies, such as dyadic social interactive behaviors or distance-based proximities.

This study aimed to determine if movement order in single-file movements are informative and effective observations for the reconstruction of animal social structures using an agent-based computational simulation model, e.g., [49, 50]. An initial artificial 2-dimensional spatial distribution was developed using the simple assumption of clustered group structures. Animal agents in a group are composed of independent and dependent agents. Independent agents distribute spatially independently each other, while dependent agents distribute depending movements of independent agents. The artificial spatial distributions aim to represent clustered structures of agent locations ‒a coupling of “core” or “keystone” subjects and “subordinate” or “follower” subjects. Collective movements were governed by two simple rules, 1) initiators of the movement are randomly chosen, and 2) the next moving agent is always the nearest neighbor of the last moving agents. These bottom-up rules allowed simulation of “single-file movement” data. Finally, we visualized social networks and reconstructed clustered structures using a recent major SNA algorithm, the Louvain algorithm, for rapid unfolding of communities in large networks [51]. The current idea ‒that artificial agents are initially generated, collective movements are produced, and collective movements are used to evaluate social networks— is similar to a previous computer experiment [52]. The current simulation is more generalized. Parameter assumptions are minimum and used mixed normal distributions. Parameter conditions were sought where single-file movement data work well for reconstruction of the latent clustered structures and further discuss possibilities for practical applications in field investigations of wild animal societies.

## Materials and Methods

The simulations described below can be used to examine whether individual-by-individual serially ordered movements, i.e., “ animal single-file movements,” provide sufficient information to reconstruct the clustered organization that is typically believed to exist in animal social groups, such as those of primates. Our simulations adopt agent-based models and involve four main steps: (1) animal agents are distributed in two-dimensional space with latent cluster organizations; (2) these generated agents are serially aligned following a simple collective movement rule (generating the “single-file movements”); (3) the social network structure is computed using an association index defined by serial orders of animal single-file movement; and (4) the cluster organization of the generated social network is evaluated. Figure 3 shows a schematic representation of the simulation process. The details of each step are described below.

### Distributing animals (1^*st*^ step)

We initially assume that the animal group of interest consists of *n*_*I*_ subgroups. Each of these subgroups has a unique *independent* animal agent (*independenter*), who moves independently of any other agent, and *n*_*D*_ *dependent* animal agents (*depender*), the spatial locations of which are invariably dependent on the location of the independenter of the subgroup. For simplicity, the number *n*_*D*_ of dependers in subgroups is assumed to be the same for all subgroups. Accordingly, the total number of animal agents is *N* = *n*_*I*_ (*n*_*D*_ + 1). We refer to the independenter of the *i*-th subgroup as the *i*-th independenter.

The locations of independenters are independently generated from the bivariate Gaussian distribution: the location of the *i*-th independenter in two-dimensional space is denoted by ***x***_*i*_ ∈ ℝ^2^ and is distributed as

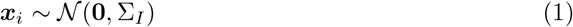

for *i* ∈ {1, ··· *n*_*I*_}. Here 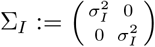 is the covariance matrix of the distribution (*σ*_*I*_ is the standard deviation of the independenter). The mean of the distribution is set to **0**, the origin of the plane.

Similarly, the locations of dependers are generated from the bivariate Gaussian distribution, but with centers dependent on the locations of independenters. The location of the *j*-th depender for the *i*-th subgroup is denoted by ***x***_*k*_ where *k* = *n*_*I*_ + (*i* × *j*) and are distributed as

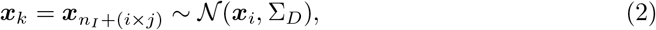

for *i* ∈ {1, ··· *n*_*I*_} and *j* ∈ {1, ··· *n*_*D*_}. Here 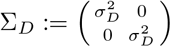 is the covariance matrix of the distribution (*σ*_*D*_ is the standard deviation of the depender). The mean parameters are set to each independenter’s locations, ***x***_*i*_. For simplicity, we always assume that *σ*_*D*_ = 1.

Consequently, the spatial distributions of *N* animal agents in the plane is dependent on three parameters, *n*_*I*_, *n*_*D*_, and *σ*_*I*_. Figure 1 shows the sample spatial distribution with the following parameter values: *n*_*I*_ = 10, *n*_*D*_ = 5, and *σ*_*I*_ = 100.0.

**Fig 1.**
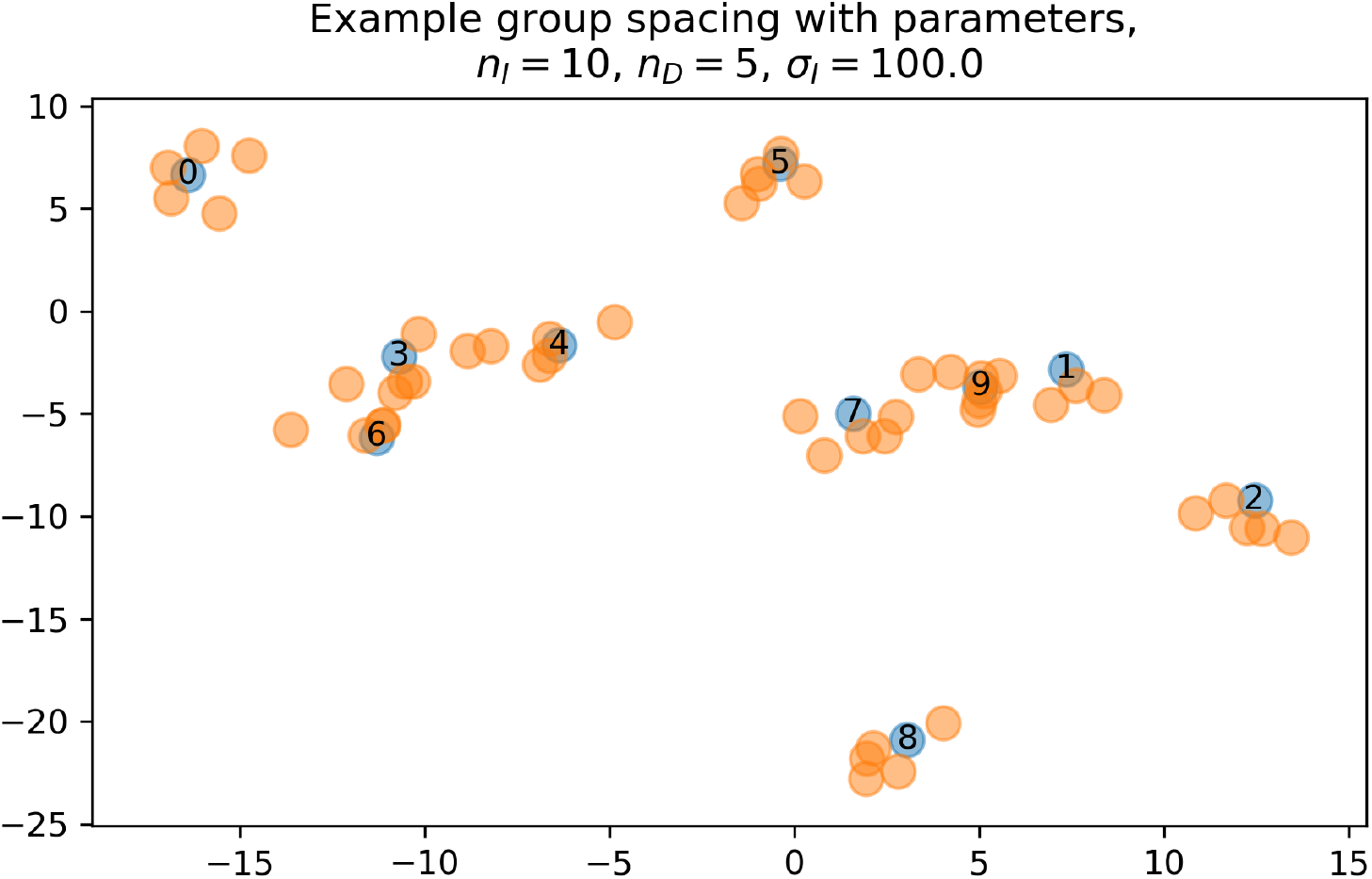
Example of the agent spatial distribution generated with model parameters, *n*_*I*_ = 10*, n*_*D*_ = 5*, σ*_*I*_ = 100.0.

### Serialization (2^*nd*^ step)

Subsequently, “animal single-file movement,” the ordered alignment set of animal agents, is generated assuming a simple rule of collective movements defined by the mutual distance between agents. The process is iterative: (1) the agent who first moves (*initiator*) is randomly selected; (2) in each iteration step, the nearest agent to the last moved agent is selected to move next. In this iterative process, the order of the single-file movement is defined as the order of movement.

More specifically, the initiator ***y***_1_ is randomly selected from a multi-set of *N* agents

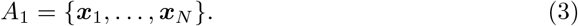

The next moving agent ***y***_2_ is selected by

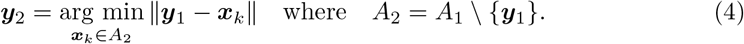

Similarly, the *i*-th moving agent is determined by

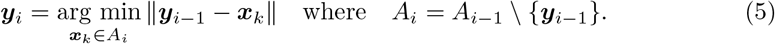

Here, || · || represents the Euclidean distance on the plane. The process of agent selection is repeated *N* − 1 times. Consequently, a totally ordered set is generated:

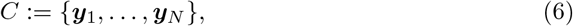

which we refer to as “animal single-file movement.” Note that the process of generating *C* depends on the three parameters, *n*_*I*_, *n*_*D*_, and *σ*_*I*_.

We repeat the process of generating the distribution of animals (1st step) and constructing the animal single-file movement (2nd step) *n*_exp_ times. Here, we implicitly refer to *n*_exp_, the number of observation opportunities for single-file movement in the field. In the following step, we therefore assume *n*_exp_ takes an experimentally feasible value, say, *n*_exp_ = 10, or 30.

Finally, we denote the set of *n*_exp_ single-file movements by 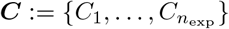.

### Generating a social network (3^*rd*^ step)

In this step, we construct a weighted graph representing the social network structure of animals using the information of ***C***.

The graph is defined by an *N × N* adjacency matrix ***G*** constructed from ***C*** as follows: First, given an animal single-file movement *C*, we define an adjacency matrix *G* as follows:

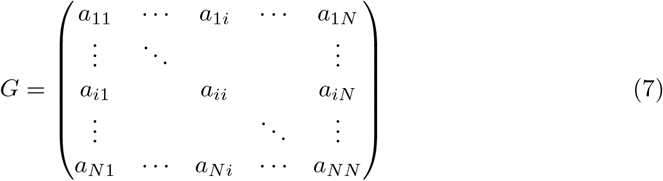

where *a*_*ij*_ = *a*_*ji*_ is 1 if the agents ***x***_*i*_ and ***x***_*j*_ are adjacent in *C*, and 0 otherwise (by definition, all diagonal elements are 0). The adjacency matrix ***G*** for ***C*** is defined as

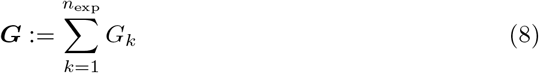

where *G*_*k*_ is the adjacency matrix for *C*_*k*_, and is the *k*-th single-file movement in ***C***.

For the implementation of this step, we use a recently widely used framework for social network analysis, `NetworkX` (https://networkx.github.io/documentation/stable/index.html) in python3 based on the from numpy matrix() method (https://networkx.github.io/documentation/stable/reference/generated/networkx.convert_matrix.from_numpy_matrix.html). We note that the *parallel edges=False* option is set, and the entries in ***G*** are interpreted as the weight of a single edge joining the vertices.

### Clustering (4^*th*^ step)

Finally, we run a graph clustering algorithm to examine whether the subgroup structure of animal agents can be reconstructed from ***G***. For this purpose, we apply the Louvain algorithm, which is one of the greedy optimization algorithms of modularity. The algorithm is implemented in the Python module `community` (or `python-louvain` on pypy) as its method `community.best partition` (see https://python-louvain.readthedocs.io/en/latest/api.html). We denote the number of clusters in the result of the Louvain algorithm by *n*_eval_. Figure 2 represents an example of a social network graph with clustering obtained using our procedure.

**Fig 2.**
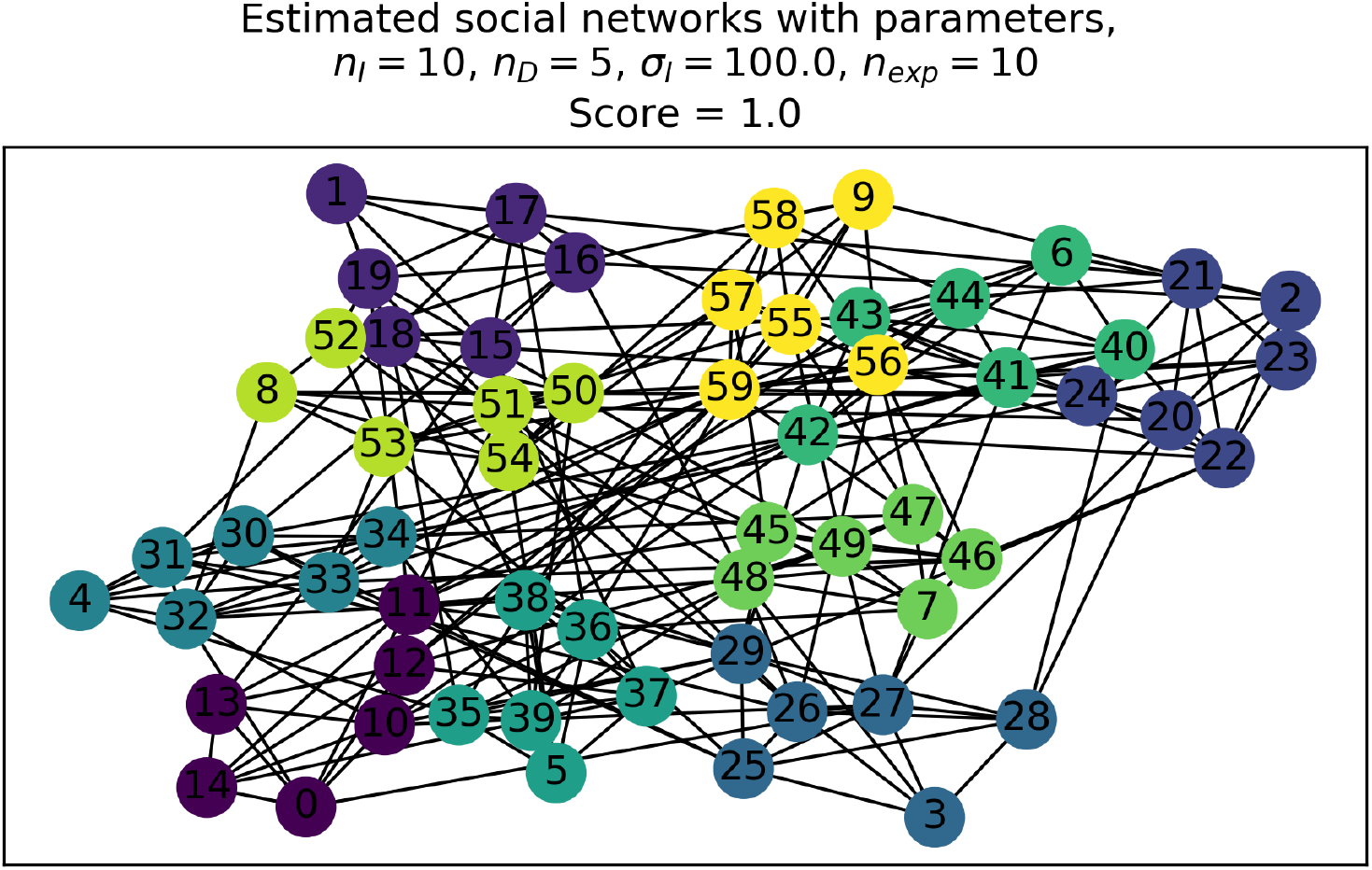
Example of a social network graph clustered using the Louvain algorithm. This graph was generated from the spacing patterns of animal agents shown in Figure 1

**Fig 3.**
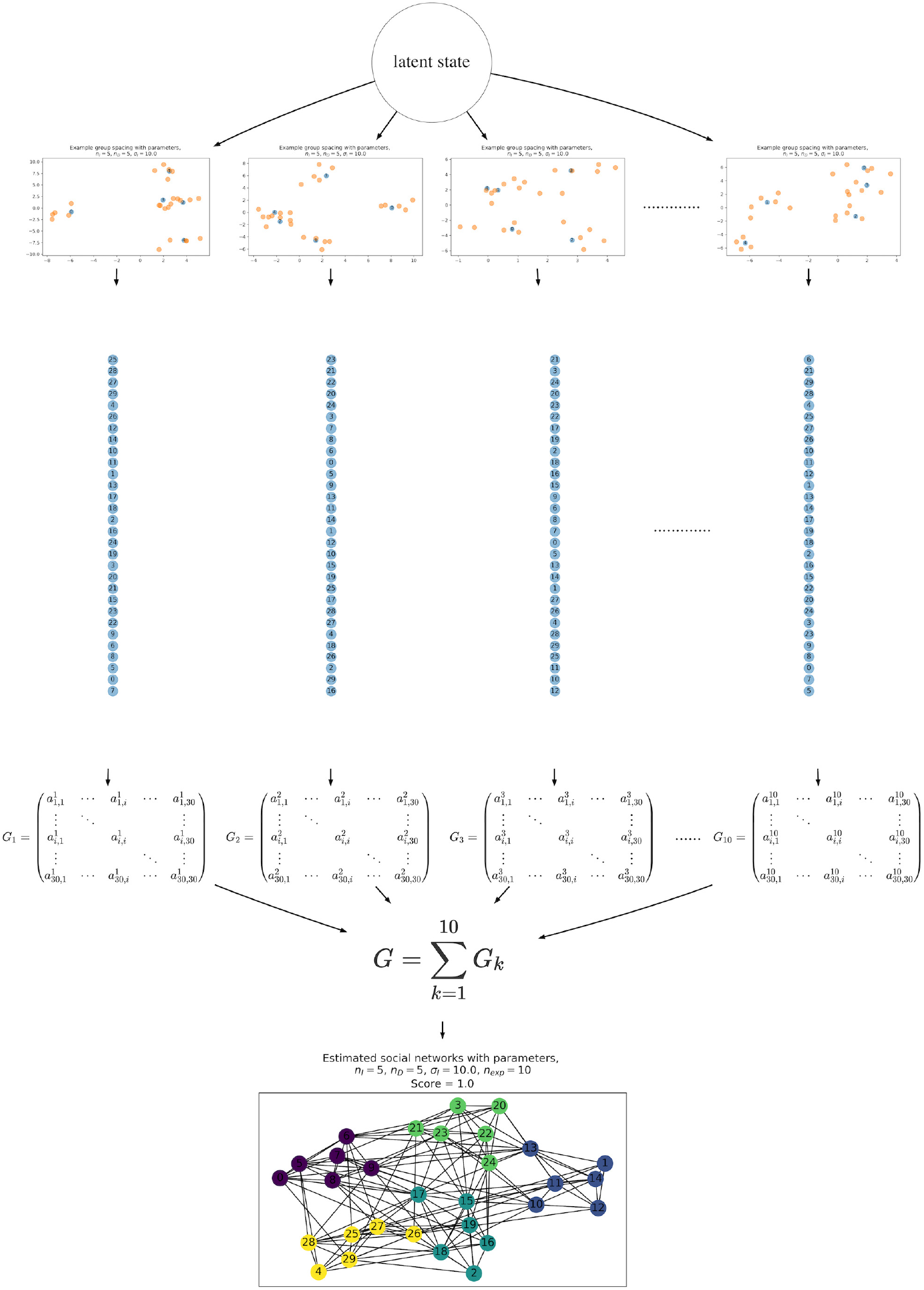
Schematic illustration of the simulation process. In this example, the parameters were set as follows: *n*_*I*_ = 5*, n*_*D*_ = 5*, σ*_*I*_ = 10*, σ*_*D*_ = 1*, n*_exp_ = 10. The simulation started from the spatial distributions of agents generated from one latent state of social group organization, determined by parameters of the mixed Gaussian processes, *n_I_, n_D_, σ_I_, σ_D_* (the two top layers). The “single-file movement” data sets were then generated using the process described in the section on ordered alignment of agents. The vertical chains of the 30 circles represent single-file movement (the number of each circle is the agent id). Adjacency matrices *G*_*k*_ for *k* ∈ {1, 2*,…*10} were generated from orders in the single-file movement data set, and were convolved to a single adjacency matrix ***G***, which was passed to the social network analysis. Finally, a social network graph was produced, with clustering of the local community (bottom graph). This flow is a single simulation process, which is run 1000 times.

### Evaluation of the method

Here, we describe our procedure for estimating the accuracy of the clustering we obtained. For this purpose, we simply count the number of clusters obtained, *n*_eval_, and compare this with the “correct answer” and the number of subgroups *n*_*I*_. Namely, we consider the ratio *r* = *n*_eval_*/n*_*I*_ and refer to this as the score of the cluster estimations. When *r* = 1, our procedure successfully recovers the information of the number of clusters, whereas *r >* 1 or *r <* 1 implies an overestimation or underestimation, respectively. For each combination of parameters *n*_*I*_ ∈ {1, 2, … 20}, *n*_*D*_ ∈ {1, 2, … 20}, *n*_exp_ ∈ {10, 30}, and *σ*_*I*_ ∈ {1.0, 2.0, 5.0, 10.0, 100.0}, we run the entire procedure 1000 times and compute the average 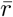 of the score *r*. The result are discussed in Section Experimental Results.

### Remarks on the method

As described in section Experimental Results, the proposed method works reasonably well for most of the selected parameter values. In this subsection, we attempt to explain why this is the case by showing that the modularity of the “true” clustering is likely to be higher than that of the clustering obtained by attaching two clusters of the true clustering together. Given that the Louvain algorithm optimizes the modularity through a process of inductive amalgamation of clusters, if the true clustering has this property, the algorithm can be expected to generate the true clustering, which implies that the score *r* will be 1.

We begin by recalling the definition of modularity. Consider a weighted graph 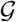 with *n* nodes defined by a symmetric *n* × *n* weighted adjacency matrix *G* = (*a*_*ij*_) and a clustering of 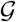 into mutually disjointed sets of nodes 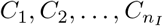 where *n*_*I*_ is the number of clusters. The modularity of this clustering is defined as

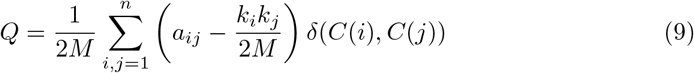

where *k*_*i*_ and *k*_*j*_ are the sum of the weights of edges from nodes *i* and *j*, respectively; *M* is the sum of the weights of all edges in the graph; *C*(*i*) and *C*(*j*) are the clusters to which nodes *i* and *j* belong, respectively, and *δ* is the Kronecker delta function. Thus, *δ*(*C*(*i*)*, C*(*j*)) is 1 if *i* and *j* belong to the same cluster but otherwise 0. Note that 2*M* = Σ_*i,j*_ *a*_*ij*_. For ***G*** construction in the 3rd step of our method, if an animal single-file movement has *N* − 1 edges, *M* = (*N* − 1) *n*_exp_ holds.

If we assume that 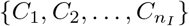 be the true clustering, that is, the clustering exactly reflects the original subgroup structure of animals, and denote its modularity by *Q*, and then consider a clustering 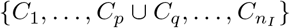 obtained from the true clustering by attaching *C*_*p*_ and *C*_*q*_ together and denote its modularity by *Q*^*′*^, a direct calculation shows that

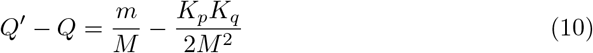

where *m* is the sum of the weights for all edges from *C*_*p*_ to *C*_*q*_; *K*_*p*_ and *K*_*q*_ are the sum of the weights for all edges from *C*_*p*_ and *C*_*q*_, respectively. We want to show that this quantity is negative; that is,

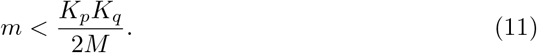

For simplicity, we assume that in each single-file movement, no mixture of subgroupsoccurs; that is, the first *n*_*D*_ + 1 agents in the movement belong to the same subgroup, and the next *n*_*D*_ + 1 belongs to another one, and so on. This is more or less equivalent to assuming that *σ*_*I*_ is very large, and therefore each each subgroup is completely separated. In this case, (11) can be reasoned as follows: A single-file movement assigns *n*_*D*_ edges to the first and last clusters and *n*_*D*_ + 1 edges to others. Therefore, *K*_*p*_, *K*_*q*_ ≤ *n*_*D*_*n*_exp_ follows. Hence, a sufficient condition for (11) to hold is as follows:

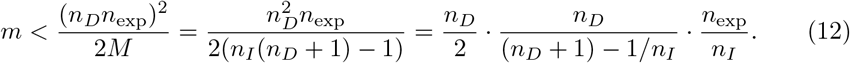

In contrast, the number of edges between agents that belong to different subgroups is *n*_*I*_ − 1 for each single-file movement. Therefore, the sum of the weights for edges in 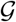 connecting different subgroups is (*n*_*I*_ − 1) *n*_exp_. Thus, the expected value of *m* is

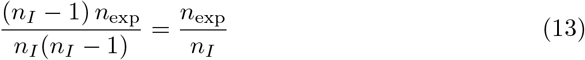

provided that all the ordering of subgroups in single-file movements is equally probable. The right-hand side of (12) can now be understood as the expected value of *m* multiplied by

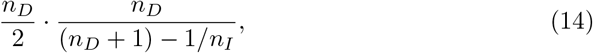

which is approximately *n*_*D*_/2 in our parameter settings. This implies that, when *n*_exp_ is sufficiently large, (12) is very likely to hold for all values of *p, q*.

## Experimental Results

Figure 4 is the contour plots of the calculated average score 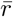 for some selected sets of parameter values. This suggests that the average score increases as parameters *σ*_*I*_ and *n*_exp_ increases, just as expected. It is also observed that the score decreases as *n*_*I*_, the numbers of the independenter, increases. In contrast, it increases as *n*_*D*_, the numbers of the depender, increases. In summary, scores will be better with large *σ*_*I*_, *n*_exp_*, n*_*D*_ and small *n*_*I*_ (expect for the case of *n*_*I*_ = 1). In the cases of *n*_exp_ = 30 and *σ*_*I*_ ≥ 5, scores were often perfect, i.e., 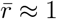. Thus, cluster size estimations were often successful, if animal single-file movements were observed 30 times.

**Fig 4.**
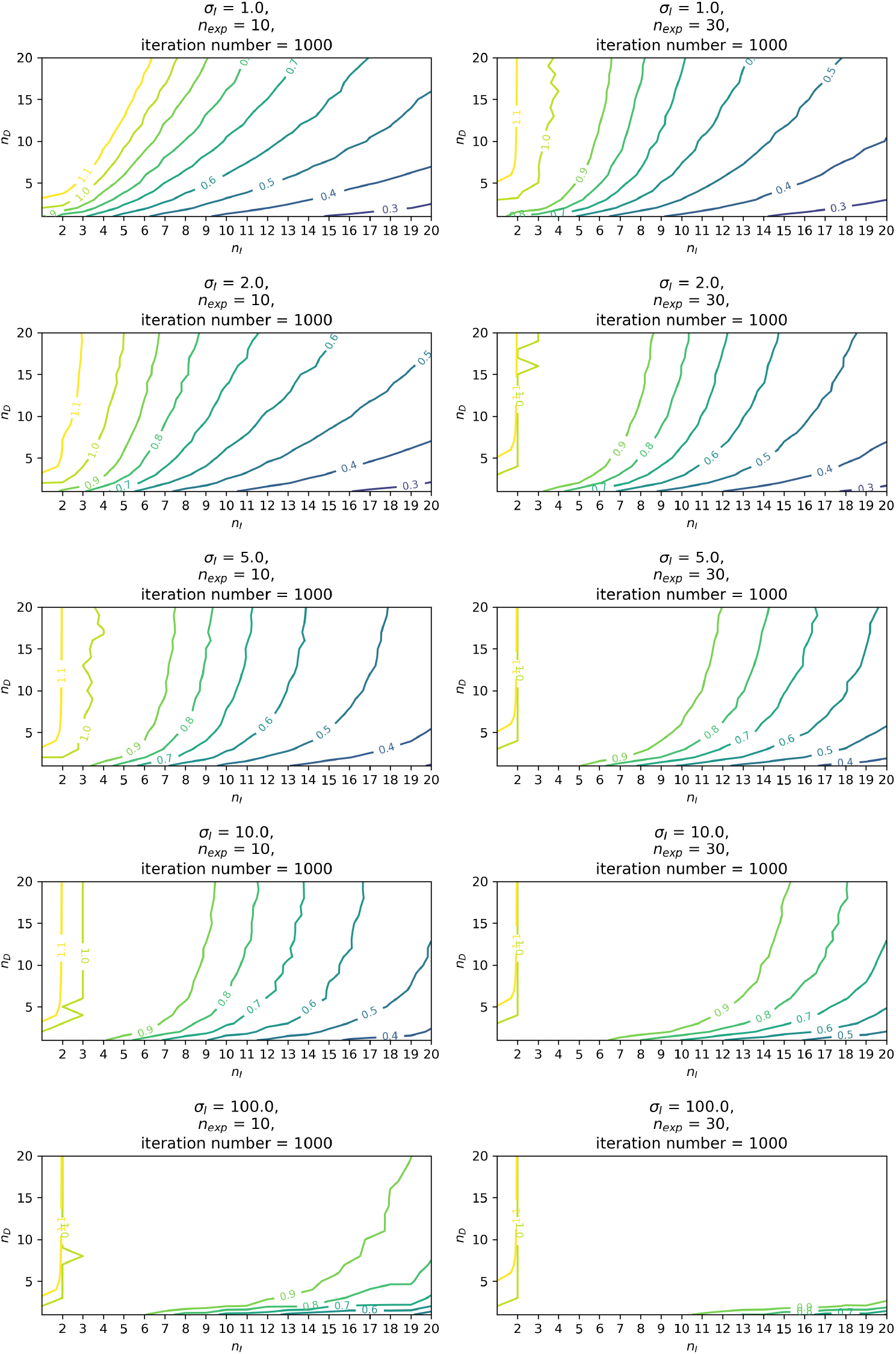
Contour plots for average performance scores for cluster estimation, 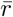 in 1000-steps simulations. Areas near the score of 1.0 represents areas of parameter combinations where perfect estimation scores returned.

## Discussion

### Result interpretations

Simulations with agent-based models suggest that recent implementation of a community clustering algorithm in the SNA could well estimate latent community structures generated by mixed Gaussian distributions under several parameter conditions. First, SD ratio parameters, 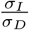, likely influence performance for identification of clustering structures. The greater these ratios are, the more precise the estimations of cluster size obtained. This result is reasonable since the contrast of the modular structure becomes clear (see Figure 1 as an example of the *σ*_*I*_ = 100). Likewise, the more iterations, the more accurate scores of clusters size became. This result is also not surprising. An important result is the effect of numbers of *dependers* for the cluster estimation. Results consistently indicate that greater numbers of *dependers* improved estimation performance, at least within the range of parameter settings used in study simulations. Too many *dependers* would likely lead to overestimations of the cluster sizes, but such confusion occurred only in the case of *σ*_*I*_ = 1*, n*_exp_ = 10, and did not occur using most other parameter combinations.

An additional lesson from the simulations is an implication that appropriate numbers of the *independenters* exist for better performance in cluster estimation. At least for parameter settings of 1 ≤ *n*_*I*_ ≤ 30, a single-digit number, except for 1 and 2, likely shows better performance. In summary, results suggested that SNA applications of the Louvain algorithm to single-file movement data work well even when using only 30 iterations in simulations (*n*_exp_ = 30), when 1) the *independenters* range between around 3 and 10, 2) *dependenters* count is as large as reasonable, and 3) SD ratios of *independenter* per *dependenter* are high.

### Practical applications: primate societies as a possible candidate

Looking back to observations of actual animal societies in reality, animals would fit with conditions appropriate for fine reconstruction using the present method? A promising candidate, for example, are primate societies.

Primates are middle to large mammals mainly inhabiting tropical forests and sometimes adapting to dry savanna, temperate forests or cool-temperate climates [53]. Primates show a large diversity of social systems [54], from simple monogamous to complex multi-level societies that presumably emerged based on the ecological constrains, such as competition for food and threats of predation, as discussed in successful ecological models, e.g., [55–57]. Their societies are the most investigated animal society from both theoretical and empirical perspectives, and the “social roles” or “social structure” have been commonly recognized under the ecological constrains, e.g., an “alpha” subject, the most dominant social rank, or the kin-based clusters, such as the female affiliative bonding [57, 58]. Thus, principles of primate social formation rely on kin-based relationships and due to ecological constrains, group size is typically limited to about 100 individuals, but mostly groups range from 2.5-50 individuals [59]. Group size and structure, which are often consist of a few matrilineal clustered families and a few independent males, seemingly match with the successful parameter conditions indicated in the simulations.

Another advantage of primates is methodological characteristics of primate behavioral ecology. Primates are typically long-lived animals with slow life histories, and long-term studies that have been ongoing continuously for approximately a decade or more, are relatively common [60, 61]. Traditionally, primatologists have made individual long-term observations. Consequently, all group members can be perfectly identified in both field and captive studies. Observations at road or river crossings ‒a suitable event for single-file movement— have been reported in field observations for several primate species, e.g., chimpanzee [62], mandrill [44] and proboscis monkeys [63]. Such events (single-file movement) would potentially happen frequently in the field, though these observations may be used only for counting group members. Relevant findings of simulations are that the SNA works well for only 30 iterations of the single-file observation, and such observations may well exist for well-studied primates. Given advantages in primate groups, applications of the present SNA to their single-file movement data can be empirically tested soon. The quest to find the social structures of the primates is a fundamental issue, studied by many pioneer primatologists. Thus, the proposed approach provides a novel and empirical tool to analyze primate society, especially for their socio-spatial organization.

### Next for the methodological generalizations

We must acknowledge that our proposed simulation approach has certain limitations. Firstly, we examined only a very limited number of parameter assumptions regarding animal social organization. For example, the number of *dependenter* per *independenter* is always constant, and the “clustering parameters,” represented by the SD ratio of 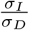, were also assumed to be constant for the *dependenters* and *independenters*. Moreover, the simulated single-file movements were observed without any misidentification of the passing agents. However, it is often difficult to reproduce the behavior of wild animals based on the strong and simple assumptions characterizing the present simulations. For methodological generalization, we must further evaluate the simulated groups of the variable sets of the parameters, or a simulated case of the single-file movement in which the identities of some agents are masked.

Ideally, the method should be validated using realistic animal data. As mentioned above, camera trapping would be a promising interface to automatically collect data on single-file movements. For primate researchers engaged in longitudinal field projects, animal identification is possible by observing video clips obtained using camera traps.

Finally, we also acknowledge possible integration with other social network data in primate societies. The output from the single-file movement-SNA is just one dimension of inference for social structures. For a deep understanding of social organizations, investigation of animal society with multidimensional views and data is needed. For example, genetic test of relatedness is a direct evaluation of network patterns. Behavioral observations, e.g., traditional quantitative grooming counts of the groomer-groomee or vocal exchanges of callers and responders, are also direct metrics that reflect social networks. Gut bacterial flora of groups of individuals was recently suggested as a possible candidate to elucidate the network structures of animal society. Thus, access to many kinds of elemental data is available for predictions concerning social networks. Together with comparisons among network structures reconstructed by different metrics, an ultimate understanding of fundamental mechanisms underlying the animal society will eventually emergence.

## Acknowledgements

We appreciate the profound comments and suggestions from Ichiro Tsuda, Aru Toyoda, and Takashi Morita. This study was performed under the Cooperative Research Program at KUPRI (2018-C-27, 2019-B-27)

## Code accessibility

All code is available from https://github.com/hkoda/mix_gauss_caravan_sna or https://doi.org/10.5281/zenodo.3876449.

## Author contributions

Project organization: HK IM; Computational simulations: HK; Analytic discussion: ZA; Manuscript writing: HK ZA IM.

## References

1. Ilany A, Akçay E. Social inheritance can explain the structure of animal social networks. Nature Communications. 2016;7(1):12084. doi:10.1038/ncomms12084.

2. Lazer D, Pentland A, Adamic L, Aral S, Barabási AL, Brewer D, et al. Social science: Computational social science. Science. 2009;323(5915):721–723. doi:10.1126/science.1167742.

3. Croft DP, James R, Krause J. Exploring animal social networks. Princeton University Press; 2008.

4. Puga-Gonzalez I, Sosa S, Sueur C. Editorial: Social networks analyses in primates, a multilevel perspective. Primates. 2019;60(3):163–165. doi:10.1007/s10329-019-00720-5.

5. Sueur C, Deneubourg JL, Petit O, Couzin ID. Group size, grooming and fission in primates: A modeling approach based on group structure. Journal of Theoretical Biology. 2011;273(1):156–166. doi:10.1016/j.jtbi.2010.12.035.

6. Krause J, James R, Franks DW, Croft DP. Animal Social Networks. Oxford University Press, USA; 2015.

7. Pasquaretta C, Levè M, Claidière N, van de Waal E, Whiten A, MacIntosh AJJ, et al. Social networks in primates: smart and tolerant species have more efficient networks. Scientific Reports. 2015;4(1):7600. doi:10.1038/srep07600.

8. Trivers RL. The Evolution of Reciprocal Altruism. The Quarterly Review of Biology. 1971;46(1):35–57. doi:10.1086/406755.

9. Silk JB, Alberts SC, Altmann J. Social Bonds of Female Baboons Enhance Infant Survival. Science. 2003;302(5648):1231–1234. doi:10.1126/science.1088580.

10. Silk JB, Beehner JC, Bergman TJ, Crockford C, Engh AL, Moscovice LR, et al. Strong and consistent social bonds enhance the longevity of female baboons. Current Biology. 2010;20(15):1359–1361. doi:10.1016/j.cub.2010.05.067.

11. Clutton-Brock T. Mammal Societies. John Wiley & Sons; 2016.

12. Clutton-Brock T, Janson C. Primate socioecology at the crossroads: Past, present, and future. Evolutionary Anthropology. 2012;21(4):136–150. doi:10.1002/evan.21316.

13. Baigger A, Perony N, Reuter M, Leinert V, Melber M, Grünberger S, et al. Bechstein’s bats maintain individual social links despite a complete reorganisation of their colony structure. Naturwissenschaften. 2013;100(9):895–898. doi:10.1007/s00114-013-1090-x.

14. Qi XG, Garber PA, Ji W, Huang ZP, Huang K, Zhang P, et al. Satellite telemetry and social modeling offer new insights into the origin of primate multilevel societies. Nature Communications. 2014;5(1):5296. doi:10.1038/ncomms6296.

15. Tavares SB, Samarra FIP, Miller PJO. A multilevel society of herring-eating killer whales indicates adaptation to prey characteristics. Behavioral Ecology. 2017;28(2):500–514. doi:10.1093/beheco/arw179.

16. Hagberg AA, Schult DA, Swart PJ. Exploring Network Structure, Dynamics, and Function using NetworkX. In: Varoquaux G, Vaught T, Millman J, editors. Proceedings of the 7th Python in Science Conference. Pasadena, CA USA; 2008. p. 11–15.

17. Farine DR, Whitehead H. Constructing, conducting and interpreting animal social network analysis. Journal of Animal Ecology. 2015;84(5):1144–1163. doi:10.1111/1365-2656.12418.

18. Kanngiesser P, Sueur C, Riedl K, Grossmann J, Call J. Grooming network cohesion and the role of individuals in a captive chimpanzee group. American Journal of Primatology. 2011;73(8):758–767. doi:10.1002/ajp.20914.

19. Levè M, Sueur C, Petit O, Matsuzawa T, Hirata S. Social grooming network in captive chimpanzees: does the wild or captive origin of group members affect sociality? Primates. 2016;57(1):73–82. doi:10.1007/s10329-015-0494-y.

20. Matsuda I, Fukaya K, Pasquaretta C, Sueur C. Factors influencing grooming social networks: insights from comparisons of colobines with different dispersal patterns. In: Dispersing Primate Females. Springer; 2015. p. 231–254.

21. Funkhouser JA, Mayhew JA, Mulcahy JB. Social network and dominance hierarchy analyses at Chimpanzee Sanctuary Northwest. PLoS ONE. 2018;13(2):e0191898. doi:10.1371/journal.pone.0191898.

22. Kawazoe T, Sosa S. Social networks predict immigration success in wild Japanese macaques. Primates. 2019;60(3):213–222. doi:10.1007/s10329-018-0702-7.

23. Lehmann J, Majolo B, McFarland R. The effects of social network position on the survival of wild Barbary macaques, Macaca sylvanus. Behavioral Ecology. 2016;27(1):20–28. doi:10.1093/beheco/arv169.

24. Castles M, Heinsohn R, Marshall HH, Lee AEG, Cowlishaw G, Carter AJ. Social networks created with different techniques are not comparable. Animal Behaviour. 2014;96:59–67. doi:10.1016/j.anbehav.2014.07.023.

25. Farine DR. Proximity as a proxy for interactions: Issues of scale in social network analysis. Animal Behaviour. 2015;104:e1–e5. doi:10.1016/j.anbehav.2014.11.019.

26. Fehlmann G, King AJ. Bio-logging. Current Biology. 2016;26(18):R830–R831. doi:10.1016/j.cub.2016.05.033.

27. Rutz C, Hays GC. New frontiers in biologging science. Biology Letters. 2009;5(3):289–292. doi:10.1098/rsbl.2009.0089.

28. Morita T, Toyoda A, Aisu S, Kaneko A, Suda-Hashimoto N, Matsuda I, et al. Animals exhibit consistent individual differences in their movement: A case study on location trajectories of Japanese macaques. Ecological Informatics. 2020;56(January):101057. doi:10.1016/j.ecoinf.2020.101057.

29. Boenisch F, Rosemann B, Wild B, Dormagen D, Wario F, Landgraf T. Tracking All Members of a Honey Bee Colony Over Their Lifetime Using Learned Models of Correspondence. Frontiers in Robotics and AI. 2018;5(APR):1–10. doi:10.3389/frobt.2018.00035.

30. Lesmerises F, Johnson CJ, St-Laurent MH. Landscape knowledge is an important driver of the fission dynamics of an alpine ungulate. Animal Behaviour. 2018;140:39–47. doi:10.1016/j.anbehav.2018.03.014.

31. Sigaud M, Merkle JA, Cherry SG, Fryxell JM, Berdahl A, Fortin D. Collective decision-making promotes fitness loss in a fusion-fission society. Ecology Letters. 2017;20(1):33–40. doi:10.1111/ele.12698.

32. Strandburg-Peshkin A, Farine DR, Couzin ID, Crofoot MC. Shared decision-making drives collective movement in wild baboons. Science. 2015;348(6241):1358–1361. doi:10.1126/science.aaa5099.

33. Westley PAH, Berdahl AM, Torney CJ, Biro D. Collective movement in ecology: from emerging technologies to conservation and management. Philosophical Transactions of the Royal Society B: Biological Sciences. 2018;373(1746):20170004. doi:10.1098/rstb.2017.0004.

34. Couzin ID, Krause J. Self-Organization and Collective Behavior in Vertebrates. In: Advances in the Study of Behavior. vol. 32. Academic Press; 2003. p. 1–75. Available from: https://linkinghub.elsevier.com/retrieve/pii/S0065345403010015.

35. Sueur C, Deneubourg JL. Self-Organization in Primates: Understanding the Rules Underlying Collective Movements. International Journal of Primatology. 2011;32(6):1413–1432. doi:10.1007/s10764-011-9520-0.

36. Peterson RO, Jacobs AK, Drummer TD, Mech LD, Smith DW. Leadership behavior in relation to dominance and reproductive status in gray wolves, Canis lupus. Canadian Journal of Zoology. 2002;80(8):1405–1412. doi:10.1139/z02-124.

37. Squires VR, Daws GT. Leadership and dominance relationships in Merino and Border Leicester sheep. Applied Animal Ethology. 1975;1(3):263–274. doi:10.1016/0304-3762(75)90019-X.

38. Mitani JC, Merriwether DA, Zhang C. Male affiliation, cooperation and kinship in wild chimpanzees. Animal Behaviour. 2000;59(4):885–893. doi:10.1006/anbe.1999.1389.

39. Fischhoff IR, Sundaresan SR, Cordingley J, Larkin HM, Sellier MJ, Rubenstein DI. Social relationships and reproductive state influence leadership roles in movements of plains zebra, Equus burchellii. Animal Behaviour. 2007;73(5):825–831. doi:10.1016/j.anbehav.2006.10.012.

40. Prins HHT. Buffalo Herd Structure and its Repercussions for Condition of Individual African Buffalo Cows. Ethology. 1989;81(1):47–71. doi:10.1111/j.1439-0310.1989.tb00757.x.

41. Perony N, Townsend SW. Why Did the Meerkat Cross the Road? Flexible Adaptation of Phylogenetically-Old Behavioural Strategies to Modern-Day Threats. PLoS ONE. 2013;8(2):e52834. doi:10.1371/journal.pone.0052834.

42. Mizuno K, Sharma N, Idani G, Sukumar R. Collective behaviour of wild Asian elephants in risky situations: how do social groups cross roads? Behaviour. 2017;154(12):1215–1237. doi:10.1163/1568539X-00003465.

43. Mitani JC, Watts DP. Correlates of territorial boundary patrol behaviour in wild chimpanzees. Animal Behaviour. 2005;70(5):1079–1086. doi:10.1016/j.anbehav.2005.02.012.

44. Hongo S. New evidence from observations of progressions of mandrills (Mandrillus sphinx): a multilevel or non-nested society? Primates. 2014;55(4):473–481. doi:10.1007/s10329-014-0438-y.

45. Aureli F, Schaffner CM, Verpooten J, Slater K, Ramos-Fernandez G. Raiding parties of male spider monkeys: Insights into human warfare? American Journal of Physical Anthropology. 2006;131(4):486–497. doi:10.1002/ajpa.20451.

46. Dore KM, Hansen MF, Klegarth AR, Fichtel C, Koch F, Springer A, et al. Review of GPS collar deployments and performance on nonhuman primates. Primates. 2020;doi:10.1007/s10329-020-00793-7.

47. Peterson RO. Wolf ecology and prey relationships on Isle Royale. 11. Department of the Interior, National Park Service; 1977.

48. O’Connell AF, Nichols JD, Karanth KU. Camera traps in animal ecology: methods and analyses. Springer Science & Business Media; 2010. Available from: https://link.springer.com/book/10.1007%2F978-4-431-99495-4.

49. Bonabeau E. Agent-based modeling: Methods and techniques for simulating human systems. Proceedings of the National Academy of Sciences. 2002;99(Supplement 3):7280–7287. doi:10.1073/pnas.082080899.

50. Grimm V. Pattern-Oriented Modeling of Agent-Based Complex Systems: Lessons from Ecology. Science. 2005;310(5750):987–991. doi:10.1126/science.1116681.

51. Blondel VD, Guillaume JL, Lambiotte R, Lefebvre E. Fast unfolding of communities in large networks. Journal of Statistical Mechanics: Theory and Experiment. 2008;2008(10):P10008. doi:10.1088/1742-5468/2008/10/P10008.

52. Sueur C, Deneubourg JL, Petit O. From Social Network (Centralized vs. Decentralized) to Collective Decision-Making (Unshared vs. Shared Consensus). PLoS ONE. 2012;7(2):e32566. doi:10.1371/journal.pone.0032566.

53. Mittermeier RA, Wilson DE, Rylands AB. Handbook of the mammals of the world: primates. Lynx Edicions; 2013.

54. Mitani JC, Call J, Kappeler PM, Palombit RA, Silk JB. The evolution of primate societies. University of Chicago Press; 2012.

55. Sterck EHM, Watts DP, Van Schaik CP. The evolution of female social relationships in nonhuman primates. Behavioral Ecology and Sociobiology. 1997;41(5):291–309. doi:10.1007/s002650050390.

56. Van Schaik CP. Why are Diurnal Primates Living in Groups? Behaviour. 1983;87(1-2):120–144. doi:10.1163/156853983X00147.

57. Wrangham RW. An Ecological Model of Female-Bonded Primate Groups. Behaviour. 1980;75(3-4):262–300. doi:10.1163/156853980X00447.

58. Shultz S, Dunbar R. Bondedness and sociality. Behaviour. 2010;147(7):775–803. doi:10.1163/000579510X501151.

59. Dunbar RIM, Carron PM, Shultz S. Primate social group sizes exhibit a regular scaling pattern with natural attractors. Biology Letters. 2018;14(1):20170490. doi:10.1098/rsbl.2017.0490.

60. Chapman CA, Corriveau A, Schoof VAM, Twinomugisha D, Valenta K. Long-term simian research sites: Significance for theory and conservation. Journal of Mammalogy. 2017;98(3):652–660. doi:10.1093/jmammal/gyw157.

61. Kappeler PM, Watts DP. Long-term field studies of primates. Springer Science & Business Media; 2012.

62. Hockings KJ. Behavioral Flexibility and Division of Roles in Chimpanzee Road-Crossing. In: Matsuzawa T, Humle T, Sugiyama Y, editors. The Chimpanzees of Bossou and Nimba. Primatology Monographs. Tokyo: Springer Japan; 2011. p. 221–229. Available from: http://link.springer.com/10.1007/978-4-431-53921-6 http://link.springer.com/10.1007/978-4-431-53921-6_24.

63. Matsuda I, Tuuga A, Akiyama Y, Higashi S. Selection of river crossing location and sleeping site by proboscis monkeys (Nasalis larvatus) in Sabah, Malaysia. American Journal of Primatology. 2008;70(11):1097–1101. doi:10.1002/ajp.20604.

